# Control of Physiologic Glucose Homeostasis via the Hypothalamic Modulation of Gluconeogenic Substrate Availability

**DOI:** 10.1101/2024.05.20.594873

**Authors:** Abdullah Hashsham, Nandan Kodur, Jiaao Su, Abigail J. Tomlinson, Warren T. Yacawych, Jon N. Flak, Kenneth T. Lewis, Lily R. Oles, Hiroyuki Mori, Nadejda Bozadjieva-Kramer, Adina F. Turcu, Ormond A. MacDougald, Martin G. Myers, Alison H. Affinati

## Abstract

The brain augments glucose production during fasting, but the mechanisms are poorly understood. Here, we show that *Cckbr*-expressing neurons in the ventromedial hypothalamic nucleus (VMN^Cckbr^ cells) prevent low blood glucose during fasting through sympathetic nervous system (SNS)-mediated augmentation of adipose tissue lipolysis and substrate release. Activating VMN^Cckbr^ neurons mobilized gluconeogenic substrates without altering glycogenolysis or gluconeogenic enzyme expression. Silencing these cells (Cckbr^TetTox^ animals) reduced fasting blood glucose, impaired lipolysis, and decreased circulating glycerol (but not other gluconeogenic substrates) despite normal insulin, counterregulatory hormones, liver glycogen, and liver gluconeogenic gene expression. Furthermore, β3-adrenergic adipose tissue stimulation in Cckbr^TetTox^ animals restored lipolysis and blood glucose. Hence, VMN^Cckbr^ neurons impact blood glucose not by controlling islet or liver physiology, but rather by mobilizing gluconeogenic substrates. These findings establish a central role for hypothalamic and SNS signaling during normal glucose homeostasis and highlight the importance of gluconeogenic substrate mobilization during physiologic fasting.

---

Mammals maintain blood glucose within a tight physiologic range by integrating environmental, nutrient, and hormonal signals across multiple organ systems. While the actions of pancreatic islet hormones on liver, adipose tissue, and muscle plays crucial roles in prandial and fasting glucose homeostasis^1^, the central nervous system (CNS) maintains blood glucose during times when glucose must be generated from stored metabolic fuels^2^, such as during inter-meal intervals (i.e., physiologic fasting) or in response to allostatic stressors such as hypoglycemia^3–6^.

Some CNS circuits, such as those in the hypothalamic arcuate nucleus (ARC), modulate islet hormone secretion and hepatic insulin sensitivity^7–10^ and others, including *Lepr*-expressing neurons of the ventromedial nucleus of the hypothalamus (VMN; VMN^Lepr^ neurons), tend to decrease blood glucose by promoting glucose uptake and metabolism^11–14^. Hence, the CNS contributes to control of glucose homeostasis at least in part by regulating islet hormone secretion and augmenting glucose disposal into metabolically active tissues.

We recently found that VMN neurons defined by expression of cholecystokinin b receptor (*Cckbr*; VMN^Cckbr^ neurons, which are largely distinct from VMN^Lepr^ neurons) contribute to the maintenance of blood glucose during physiologic fasting, in addition to driving glucose production during the counter-regulatory response (CRR) to hypoglycemia^15^. In contrast to previously described neuronal populations, however, silencing VMN^Cckbr^ neurons lowers blood glucose during physiologic fasting without altering islet hormone secretion or glucose disposal, but rather by controlling glucose production^15^. Thus, VMN^Cckbr^ neurons must control blood glucose by modulating sympathoadrenal (i.e., epinephrine/norepinephrine and glucocorticoid) function, liver gluconeogenic capacity, and/or the liberation of gluconeogenic substrates from tissue storage depots. We previously showed that VMN^Cckbr^ neuron activation augments sympathoadrenal output, but it remains unclear whether these neurons also alter gluconeogenic capacity and/or other aspects of liver physiology.

Here, we identify the previously unknown mechanisms by which VMN^Cckbr^ neurons control glucose homeostasis during physiologic fasting. The acute activation of VMN^Cckbr^ neurons, as during allostatic challenges like hypoglycemia, increases sympathoadrenal output and promotes the mobilization of several gluconeogenic substrates to rapidly increase blood glucose without altering islet or liver physiology. Similarly, VMN^Cckbr^ neurons maintain fasting glucose concentrations without altering circulating hormones or liver physiology, but instead support sympathetic outflow to white adipose tissue (WAT), promoting lipolysis to provide a continuous source of glycerol to fuel glucose production. These data reveal a previously unrecognized islet hormone- and glucose uptake-independent gluconeogenic substrate-driven mechanism by which the CNS controls fasting blood glucose.

## Results

### Activating VMN^Cckbr^ neurons mobilizes gluconeogenic and ketogenic substrates

To understand the mechanisms by which VMN^Cckbr^ neuron activation increases blood glucose, we injected AAV^DIO-ChR2-GFP^ into the VMN of *Cckbr^Cre^* mice and placed a fiber optic cannula above the injection site (Cckbr^ChR2^ animals), permitting the activation of VMN^Cckbr^ cells by blue light. Consistent with our previous findings, optogenetic activation of VMN^Cckbr^ neurons rapidly mobilized glucose (Fig 1A); this stimulus produced no change in liver glycogen content after 30 minutes, however (Fig 1B). Furthermore, although pretreatment of the Cckbr^ChR2^ animals with the glycogenolysis inhibitor CP91,149 tended to decrease blood glucose in the absence of optogenetic stimulation as previously reported^16^, this treatment did not interfere with robust glucose mobilization in response to blue light (Fig 1C). Hence, liver glycogenolysis contributes minimally, if at all, to glucose mobilization during the activation of VMN^Cckbr^ neurons.

**Figure 1:**
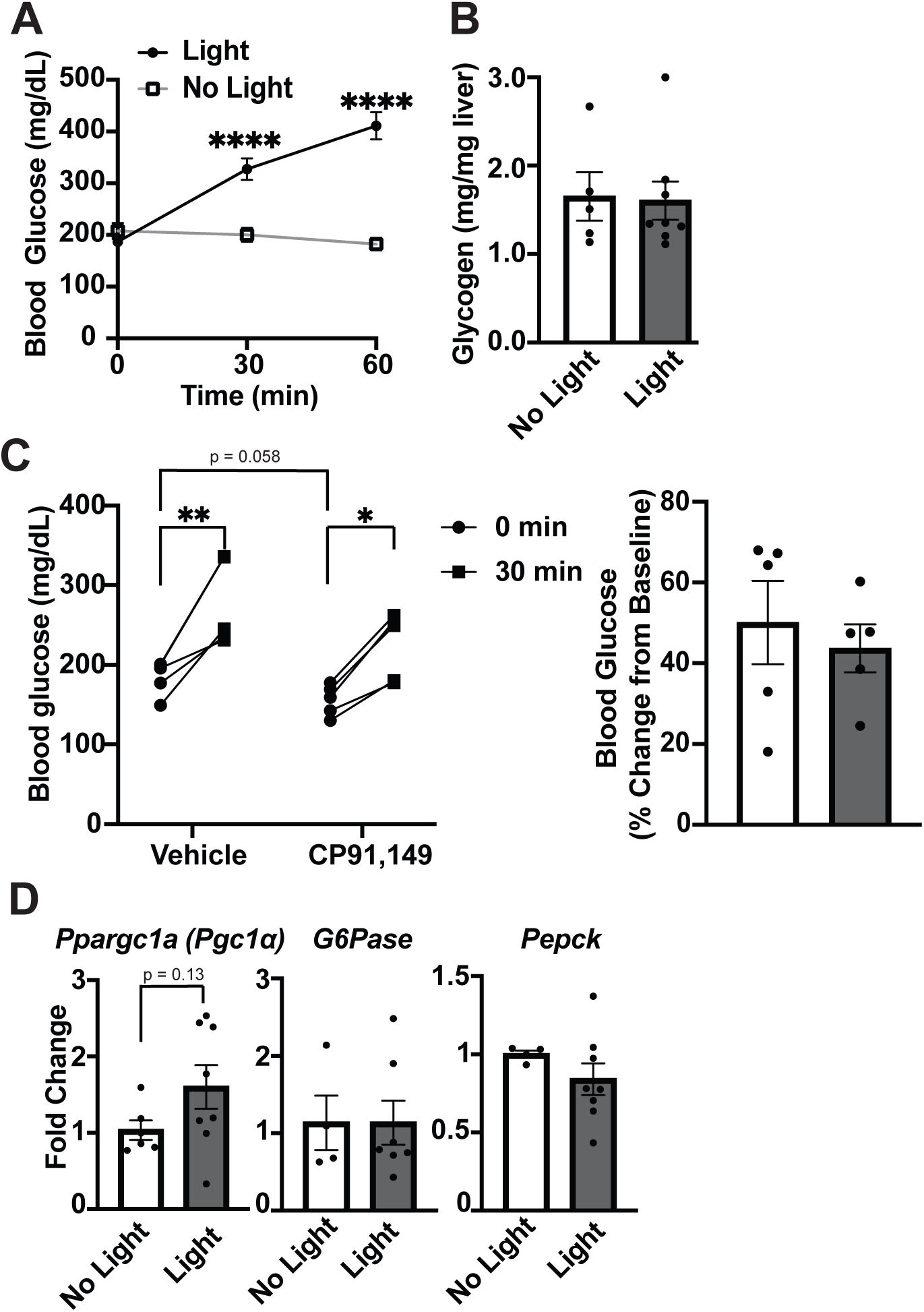
Activating VMN^Cckbr^ neurons mobilizes glucose without altering glycogen or gluconeogenic gene expression. Cckbr^Cre^ mice were injected with AAV-DIO-ChR2-eYFP unliterally into the VMN followed by optogenetic fiber placement to permit optogenetic activation of VMN^Cckbr^ neurons. Mice were fasted for 2 hours during the early light cycle prior to testing. Shown is **(A)** the glycemic response to light or no light control stimulation (n = 19) and **(B)** liver glycogen content from tissue collected after 30 minutes of light or control stimulation (n = 5 no light, 8 light). **(C)** A separate cohort of mice were treated with CP91,149, a glycogenolysis inhibitor, or vehicle at the start of the fasting period and glucose was measured before and 30 minutes after optogenetic stimulation (n = 5); the right-hand panel shows the percent change for light-stimulated vs no light conditions for each treatment. **(D)** mRNA expression of hepatic gluconeogenic genes was quantified following 30 minutes control or light delivery (n = 6 light off, 8 light on). Data are plotted as mean SEM. *p< 0.05, **p< 0.01, ****p<0.0001, by paired Student’s t-test (A, C) and unpaired Student’s t-test (B, C(t=0 comparison), D).

We also measured the expression of key gluconeogenic genes in the liver following the activation of VMN^Cckbr^ neurons (Fig 1D). Consistent with the rapid onset and peak of glucose mobilization during VMN^Cckbr^ neuron activation (before transcriptionally mediated changes could take effect), the expression of *Ppargc1a (Pgc1α)* and the gluconeogenic genes *Pck* (*Pepck)* and *G6pc* (*G6pase)* were unchanged by optogenetic stimulation (Fig 1D).

We previously found that activating VMN^Cckbr^ neurons, like the CRR, promotes the release of corticosterone and catecholamines without altering islet hormones^15^. Because these counterregulatory mediators act on liver, muscle, and fat to promote the release of gluconeogenic substrates, we measured plasma pyruvate, lactate, amino acids, and glycerol following the optogenetic stimulation of VMN^Cckbr^ neurons. We found that VMN^Cckbr^ neuron activation increased circulating pyruvate, lactate, and glycerol, although amino acids did not change (Fig 2A).

**Figure 2:**
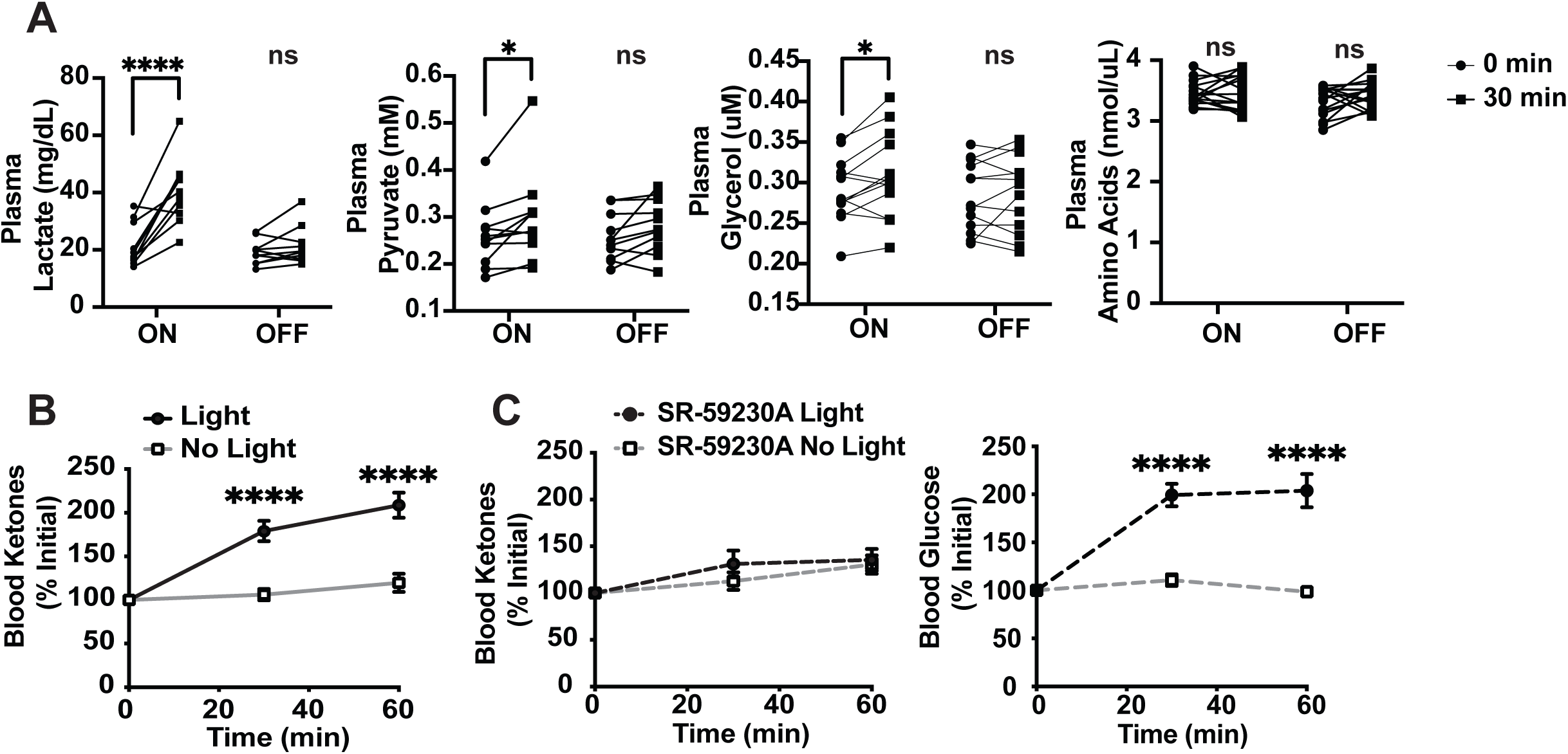
VMN^Cckbr^ neuron activation mobilizes gluconeogenic substrates and ketones. Plasma concentration of gluconeogenic substrates were measured following 30 minutes optogenetic stimulation of VMN^Cckbr^ neurons and with no light control. Shown are concentrations of **(A)** plasma lactate, pyruvate, glycerol and amino acids (n = 10-15) and **(B)** and blood ketones (n = 9). **(C-D)** Blood ketone and glucose concentrations were measured at 30 minutes intervals during light stimulation in mice pre-treated with β3-adrenergic receptor antagonist SR-59230A or vehicle (n = 9). Data are plotted as individuals **(A)** or mean +/- SEM **(B-D)** and analyzed by paired Student’s t-test. *p< 0.05, ****p<0.0001.

VMN^Cckbr^ neuron signaling is necessary for appropriate glucose mobilization during starvation, insulinopenic diabetes, and CRR-conditions in which the utilization of lipid stores provides an alternative source of energy via ketone bodies^15,17^. We thus predicted that activating VMN^Cckbr^ neurons would increase ketone body production. Indeed, VMN^Cckbr^ neuron stimulation robustly increased blood ketones (Fig 2B). Treating mice with a β3-adrenergic receptor antagonist (SR-59230A), which primarily blocks adrenergic signaling to adipose tissue, abrogated ketone production but not glucose mobilization/hyperglycemia (Fig 2C). These findings demonstrate that VMN^Cckbr^ neuron activation promotes ketone production and that this ketogenesis (but not the accompanying glucose production) requires adipose tissue lipolysis, consistent with the idea that fatty acids are needed for ketone production but that gluconeogenic substrates other than glycerol (e.g., pyruvate, lactate, and/or amino acids) that derive from non-adipose tissue sources suffice to support glucose production in response to optogenetic VMN^Cckbr^ neuron activation.

### Silencing VMN^Cckbr^ neurons impairs gluconeogenic substrate delivery from adipose tissues

Tetanus toxin-mediated silencing of VMN^Cckbr^ neurons (Cckbr^TetTox^ mice) reduces blood glucose across the diurnal cycle, with the largest difference occurring at the onset of the light period, when food intake rapidly drops (Fig 3A). This suggests that the activity of these neurons contributes not only to the response to allostatic stimuli^15^, but also to the homeostatic control of blood glucose during the early stages of fasting. We thus sought to establish the mechanisms by which physiologic action of VMN^Cckbr^ neurons modulates fasting blood glucose.

**Figure 3:**
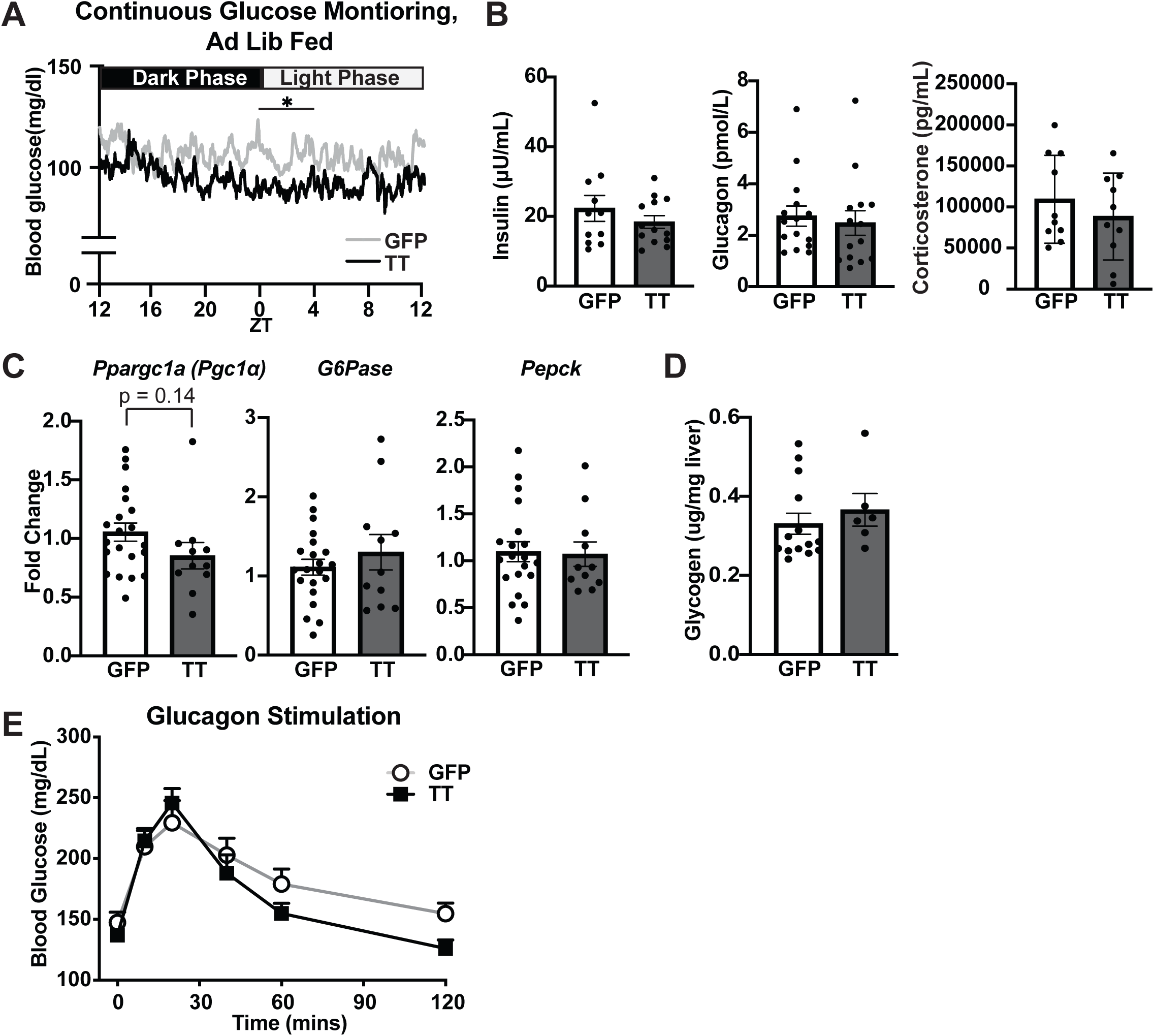
Silencing VMN^Cckbr^ neurons decreases glucose during physiologic fasting without altering pancreatic hormones, glycogen, or gluconeogenic gene expression. We injected Cckbr^Cre^ mice with AAV-DIO-TetTox-EGFP bilaterally into the VMN (n = 11) or AAV-DIO-GFP as a control (n = 10). **(A)** Blood glucose over a 24 hour period was monitored via continuous glucose monitoring in a subset of mice (n = 4 Cckbr^TetTox^ and 5 control)(ZT = Zeitgeber Time, TT = Tetanus Toxin). Following a 4 hour fast we measured **(B)** plasma hormones, **(C)** hepatic gluconeogenic gene expression, and **(D)** liver glycogen content. **(E)** Mice were injected with glucagon following a 4 hour fast and blood glucose was measured (n = 15 Cckbr^TetTox^ and 10 control). Data are plotted as mean +/- SEM, analyzed by unpaired Student’s t-test. Differences in blood glucose by CGM was analyzed in 4-hr circadian windows by 2-way ANOVA. *p<0.05

We measured islet and adrenal hormones following a 4-hour fast during the early light phase (Fig 3B), revealing unchanged insulin, glucagon, and corticosterone in Cckbr^TetTox^ compared to control mice (despite Cckbr^TetTox^ animals having lower blood glucose levels). The finding of unchanged islet hormones in Cckbr^TetTox^ animals is consistent with the failure of optogenetically stimulating VMN^Cckbr^ neurons to alter insulin or glucagon^15^. Although VMN^Cckbr^ neuron activation increases corticosterone^15^, the normal circulating corticosterone of Cckbr^TetTox^ animals suggests that processes other than those mediated by VMN^Cckbr^ neurons suffice to maintain baseline corticosterone concentrations during physiologic fasting.

Despite the chronic nature of VMN^Cckbr^ neuron silencing in Cckbr^TetTox^ animals, which should provide adequate opportunity for any alterations in gene expression to manifest, the expression of gluconeogenic genes in the liver remained unchanged in Cckbr^TetTox^ animals compared to controls (Fig 3C), as did liver glycogen content (Fig 3D). Thus, the decreased blood glucose of Cckbr^TetTox^ animals during physiologic fasting does not result from changes in circulating hormones, liver glycogen, or gluconeogenic gene expression.

To determine whether decreased blood glucose in Cckbr^TetTox^ mice might result from impaired sensitivity to glucagon and/or the inability to mobilize glycogen stores, we monitored blood glucose in Cckbr^TetTox^ mice and controls following the injection of glucagon, which rapidly mobilizes hepatic glycogen and increases hepatic gluconeogenic capacity over the longer term. We observed similarly robust initial glucose excursions in Cckbr^TetTox^ mice and controls following glucagon injection, although blood glucose levels decreased more rapidly in Cckbr^TetTox^ mice (Fig 3E). These data are consistent with normal short-term glucagon-stimulated glucose mobilization in Cckbr^TetTox^ mice, despite a potentially impaired long-term gluconeogenic response.

Overall, these findings suggest that the low blood glucose of Cckbr^TetTox^ mice does not result from alterations in islet hormones, corticosterone, or the glycogenolytic response to glucagon. Furthermore, any alteration in gluconeogenesis in the Cckbr^TetTox^ animals does not arise from changes in liver gluconeogenic gene expression. Hence, given the robust mobilization of gluconeogenic substrates by VMN^Cckbr^ neuron activation, we tested the hypothesis that low gluconeogenic substrate availability underlies the low fasting blood glucose observed in Cckbr^TetTox^ animals.

We measured plasma concentrations of lactate, pyruvate, amino acids, and glycerol in Cckbr^TetTox^ animals following a 4-hour fast. While lactate, pyruvate, and amino acids were not different between Cckbr^TetTox^ mice and controls, Cckbr^TetTox^ mice exhibited plasma glycerol levels 38% lower than those in control mice (Fig 4A). To determine whether this alteration in glycerol might reflect a decrease in adipose tissue lipolysis, we also examined plasma nonesterified fatty acids (NEFA, Fig 4B), revealing decreased NEFA in Cckbr^TetTox^ mice.

**Figure 4:**
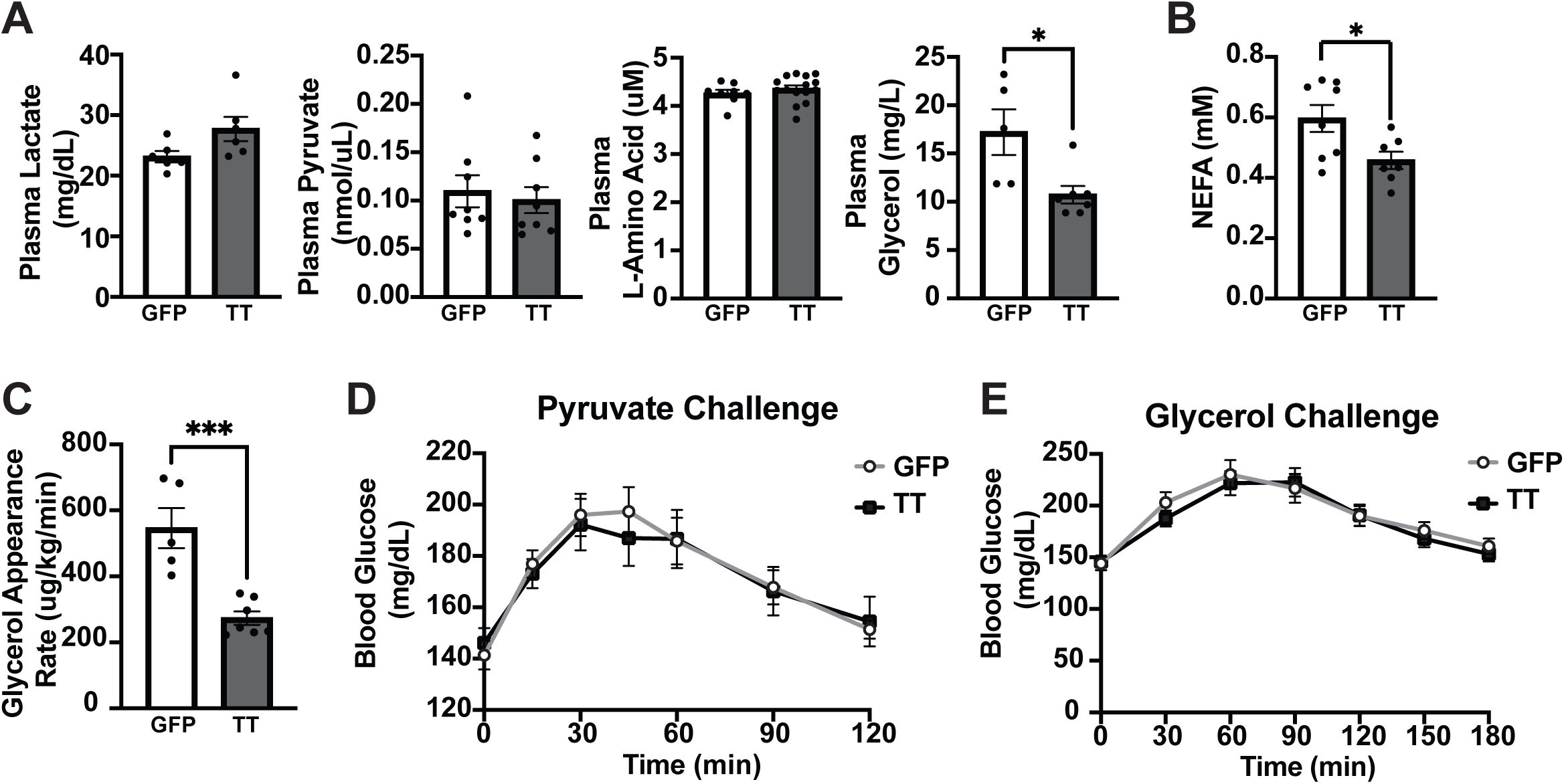
Silencing VMN^Cckbr^ neurons impairs glycerol mobilization and lipolysis. (A) Gluconeogenic substrates were measured following a 4 hour fast in Ccbkr^TetTox^ (n = 7) and GFP control mice (n = 5). **(B)** Plasma NEFA concentration was measured following a 4 hour fast. **(C)** Animals underwent a glycerol appearance assay with infusion of [^3^H]-labelled glycerol and glycerol appearance rate was calculated. Ccbkr^TetTox^ and control mice were injected with **(D)** pyruvate (2g/kg) or **(E)** glycerol (2 g/kg) following a 4-hour fast and blood glucose was measured at the indicated times. Data are plotted as mean +/- SEM. *p < 0.05, ***p < 0.001, by unpaired Student’s t-test.

Because alterations in production or utilization can each impact the concentration of circulating metabolites, we infused radiolabeled glycerol in these mice to directly examine the rate of glycerol appearance in the circulation^18^, revealing decreased glycerol production in the Cckbr^TetTox^ mice compared to controls (Fig 4C). While decreased adipose tissue lipolysis could theoretically decrease blood glucose by increasing glucose oxidation (e.g., in skeletal muscle) to compensate for lost fatty acid oxidation, we previously showed that VMN^Cckbr^ neurons promote glucose production without altering glucose disposal/utilization^15^. Hence, these findings suggest that lower baseline adipose tissue lipolysis in Cckbr^TetTox^ mice may reduce blood glucose by decreasing glycerol-mediated gluconeogenesis.

If the impairment in glucose production in Cckbr^TetTox^ mice results from limitations in gluconeogenic substrate delivery in the face of otherwise normal liver gluconeogenic capacity, Cckbr^TetTox^ mice should exhibit normal glycemic responses to gluconeogenic substrate infusion. We thus performed pyruvate (Fig 4D) and glycerol (Fig 4E) challenges in Cckbr^TetTox^ mice, revealing similar glucose excursions in Cckbr^TetTox^ mice and controls in both cases. Thus, Cckbr^TetTox^ mice exhibit normal hepatic gluconeogenic responses to substrate infusion, including in response to glycerol. Together, these findings led us to test whether the decreased blood glucose in Cckbr^TetTox^ mice results from decreased gluconeogenesis due to limitations in substrate availability that stem from impaired adipose tissue lipolysis.

### Impaired SNS outflow to WAT decreases lipolysis and glycerol-dependent gluconeogenesis in Cckbr^TetTox^ mice

SNS outflow to WAT drives lipolysis through activation of β3-adrenergic receptors, expression of which is predominantly restricted to adipose tissue in mice^19–21^. To determine whether decreased SNS outflow to WAT might underlie the decreased lipolysis and low circulating glycerol observed during physiologic fasting in Cckbr^TetTox^ mice, we treated animals with the β3-adrenergic receptor agonist CL-316,243 to stimulate β3-adrenergic receptors, bypassing the neural/SNS control of lipolysis. While Cckbr^TetTox^ mice exhibited decreased plasma glycerol and *in vivo* lipolysis following a 4-hour fast, CL-316,243 robustly increased both glycerol and lipolysis in these animals (Fig 5A-B). The resulting CL-316,243-stimulated plasma glycerol concentrations and appearance rates did not differ between control and Cckbr^TetTox^ mice (Fig 5A-B). Hence, direct SNS stimulation of adipose tissue equalizes lipolysis and glycerol availability in Cckbr^TetTox^ and control mice.

**Figure 5:**
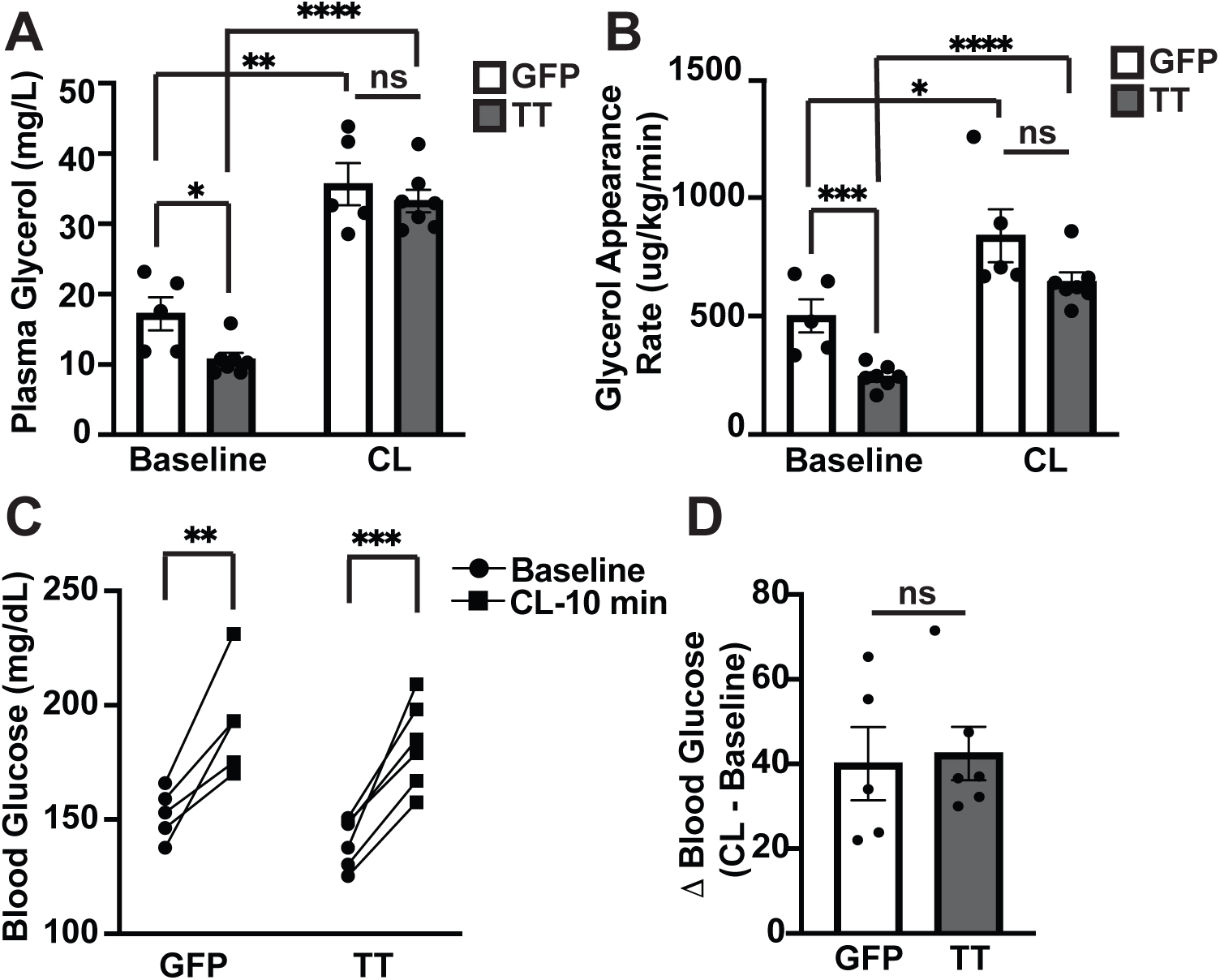
Restoring sympathetic signaling ameliorates impaired lipolysis and glucose mobilization during physiologic fasting in Cckbr^TetTox^ mice. (A) Glycerol, **(B)** glycerol appearance rate, and **(C-D)** blood glucose were measured in Cckbr^TetTox^ and control mice following a 4 hour fast at 0 and 10 minutes after β3-agonist CL316,243 i.v. infusion (n = 5). Data are plotted as mean SEM. *p < 0.05, ***p < 0.001, analyzed by unpaired Student’s t-test.

Because increasing SNS activation of WAT *in vivo* increases lipolysis and plasma glycerol concentrations, we predicted that this increase in gluconeogenic substrate availability would also increase glucose production. Indeed, treating mice with CL-316,243 similarly increased blood glucose in Cckbr^TetTox^ and control animals (Fig 5C-D). Hence, restoring adipose tissue lipolysis alleviates the glucose production defect caused by silencing VMN^Cckbr^ neurons. This suggests that VMN^Cckbr^ neurons regulate SNS outflow to WAT, promoting lipolysis to provide glycerol that fuels the gluconeogenesis needed to maintain normoglycemia during in the early stages of physiologic fasting.

To understand the cause of decreased adipose tissue lipolysis in Cckbr^TetTox^ mice, we collected and analyzed gonadal and subcutaneous WAT (gWAT and psWAT, respectively) depots from Cckbr^TetTox^ mice 10 – 12 weeks after viral injection. While we found no significant difference in adipose depot mass or adipocyte size between Cckbr^TetTox^ and control GFP mice in either WAT depot (Fig 6A-B), adipose depot mass tended to be increased in Cckbr^TetTox^ mice. As Cckbr^TetTox^ WAT depot size is not lower than controls, the observed decrease in lipolysis in these animals does not result from decreased lipid stores. The variance in adipocyte size differed significantly between Cckbr^TetTox^ mice and controls for both depots (F = 8.29, DFn = 7, DFd = 5, p = 0.03 for gWAT and F = 18.39, DFn = 7, DFd = 5, p = 0.005 for psWAT) (Fig 6A-B), however, consistent with altered adipose tissue remodeling due to impaired lipolysis in the Cckbr^TetTox^ adipose tissue^22^. This finding, in combination with our observation that pharmacologic activation of SNS signaling restores WAT lipolysis in Ccbkr^TetTox^ mice suggests that the reduction in glycerol and lipolysis results from impaired SNS outflow to WAT, rather than an intrinsic defect in WAT function.

**Figure 6:**
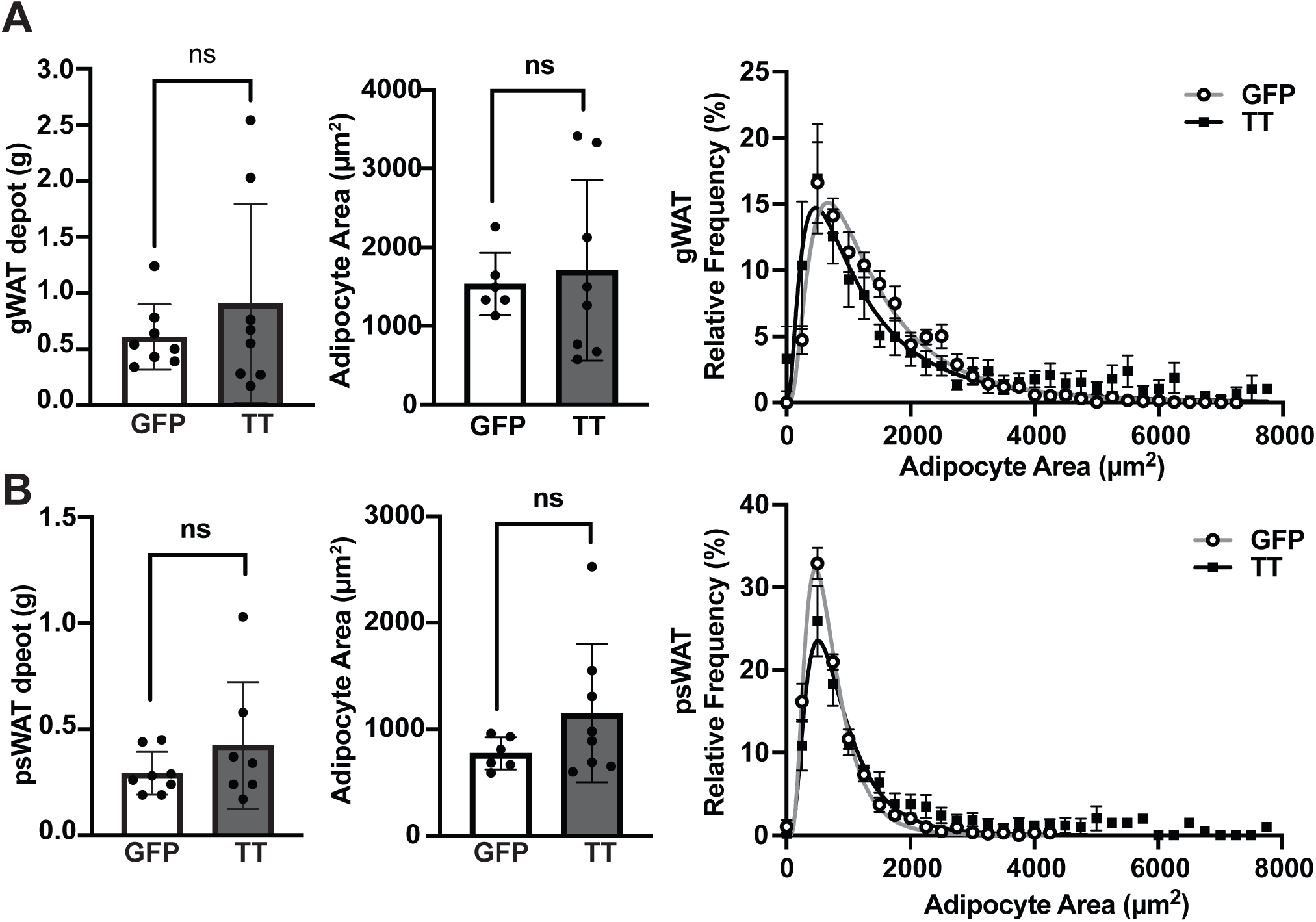
Silencing VMN^Cckbr^ neurons does not decrease adipose depots. We injected AAV-DIO-TetTox-EGFP and GFP control virus bilaterally into the VMN of Cckbr^Cre^ mice. At least 8 weeks after surgery, we collected gonadal and posterior subcutaneous white adipose tissue, (gWAT and psWAT respectively). Shown are **(A)** gWAT and **(B)** psWAT adipose tissue depot mass, average adipocyte area and adipocyte area versus frequency. (n = 6 - 8). Data are plotted as mean +/- SEM.

## Discussion

Our findings demonstrate that VMN^Cckbr^ neurons do not alter glycogenolysis, gluconeogenic gene expression, or pancreatic hormones, but rather maintain appropriate blood glucose by mobilizing gluconeogenic substrates. While artificial activation of VMN^Cckbr^ cells increases the availability of multiple gluconeogenic substrates (as during allostatic responses), during the early stages of physiologic fasting VMN^Cckbr^ neurons mainly promote SNS-mediated WAT lipolysis to provide glycerol as a gluconeogenic substrate. In addition to revealing that CNS-directed control of gluconeogenic substrate mobilization plays a crucial role in determining blood glucose concentrations during fasting, these findings demonstrate that distinct CNS circuits control the individual processes (e.g., pancreatic hormone secretion, liver physiology, and substrate mobilization) that contribute to glucose production and fasting glucose homeostasis.

The CNS directs many processes that regulate glucose during allostatic responses^14,15,23–25^. Indeed, global VMN activation increases blood glucose by modulating pancreatic hormone secretion^25^, glycogenolysis, gluconeogenic gene expression^25^, and sympathoadrenal function^15,25^. The CNS also controls glucose uptake and utilization by brown adipose tissue (BAT) and skeletal muscle, including via the VMN^Lepr^ subset of VMN neurons, which is largely distinct from VMN^Cckbr^ cells)^11–15^.

In contrast, we found that activating the VMN^Cckbr^ subset of VMN neurons promotes glucose production primarily via the sympathoadrenal mobilization of gluconeogenic substrates. Hence the mobilization of gluconeogenic substrates by the CNS suffices to promote glucose production, and this process is separable from other aspects of CNS-mediated glucose mobilization during allostatic-like responses. While activating VMN^Cckbr^ neurons increases glycerol, pyruvate, and lactate, it does not alter circulating amino acid concentrations. This suggests either that long-term activation is required to mobilize amino acids or that amino acid conversion to glucose is sufficiently rapid that our methods are unable to detect underlying changes in amino acid mobilization. Alternatively, a distinct circuit may regulate protein degradation and amino acid mobilization.

Globally interfering with VMN function (e.g., by deleting VMN vGLUT2, thereby blocking glutamatergic signaling by VMN neurons) mediates effects opposite to those of VMN activation, impairing glucagon secretion, glycogenolysis, and hepatic transcriptional responses to hypoglycemia and during fasting^26^. While silencing the VMN^Cckbr^ subset of VMN neurons decreases fasting blood glucose, it does not alter pancreatic hormone secretion, glycogen responsiveness/glycogenolysis, or long-term hepatic gluconeogenic gene expression. Rather, Cckbr^TetTox^ mice exhibit decreased glycerol production and a normal hyperglycemic response to exogenous substrates, suggesting that within the VMN, distinct cell types may control different components of the glycemic response to fasting.

While hepatic gluconeogenesis uses multiple substrates (e.g., amino acids and TCA cycle-derived lactate and pyruvate, in addition to glycerol), hormonal signaling and the availability of individual substrates dictates the relative importance of each under a distinct conditions. Our data demonstrate that during early physiologic fasting, glycerol represents a key gluconeogenic substrate for the maintenance of adequate blood glucose. Indeed, other recent studies have demonstrated the importance of glycerol for hepatic gluconeogenesis under physiologic conditions^27–29^.

The VMN controls sympathetic outflow to multiple metabolic target organs, including islets^8^, liver, and BAT^11^. Our finding that VMN^Cckbr^ neurons support lipolysis are consistent with previous reports that manipulating VMN neuronal activity alters FFA mobilization and lipid metabolism^30–32^. However, to date, retrograde, polysynaptic viral-based tracing studies from WAT have identified few VMN neurons (which appear only after prolonged infection periods^33^). While other hypothalamic sites (such as the PVN and DMH, which lie downstream from the VMN) connect more directly to WAT, VMN^Cckbr^ neurons do not project to these regions^15,34^. Thus, the downstream neuroanatomic pathways through which VMN^Cckbr^ neurons mediate WAT lipolysis remain to be defined.

Integrating these findings into a comprehensive model of overall glucose homeostasis, VMN^Cckbr^ neurons (and the CNS in general) play a crucial role in maintaining adequate blood glucose during physiologic fasting and allostatic responses, while CNS effects are less prominent during feeding. As insulin decreases during fasting and hormonal and nutrient signals drive liver physiology toward gluconeogenesis, the brain increases lipolysis through activation of the SNS to provide glycerol for gluconeogenesis, while continued low level insulin action prevents unrestrained catabolism in WAT and skeletal muscle.

In response to strong allostatic challenges, such as hypoglycemia, the CNS coordinates the CRR to promote glucagon release and suppress insulin secretion, which augments glycogenolysis and gluconeogenic capacity. Simultaneously, adrenal glucocorticoid secretion and direct tissue SNS output mobilize gluconeogenic substrates to prevent low blood glucose^35,36^. Whereas direct glucose sensing by glucagon-secreting islet alpha cells contribute to glucagon secretion, intact CNS circuit signaling (including through the VMN)^1,25,26^ is necessary for adequate glucose mobilization. VMN^Cckbr^ neurons contribute to the sympathoadrenal responses that provide the substrate to drive gluconeogenesis (and ketogenesis) during allostatic responses, but do not control glucagon secretion^15^; thus, additional circuits that contribute to other aspects of the CRR remain to be identified.

In type 1 diabetes, the absolute lack of insulin increases hepatic gluconeogenic capacity and derepresses lipolysis and proteolysis, promoting unrestrained release of substrates for gluconeogenesis (and ketogenesis) and raising blood glucose. Because silencing VMN^Cckbr^ neurons decreases blood glucose in models of type 1 diabetes^15^, CNS-mediated processes also contribute to the substrate mobilization that contributes to the hyperglycemia of insulinopenic diabetes.

The signals that modulate VMN^Cckbr^ neuronal activity (and other CNS neurons that control glucose production) during physiologic fasting, allostatic responses, and insulinopenic diabetes remain unknown and thus represent important future research directions. To understand how dysregulation of VMN^Cckbr^ neurons and similar CNS systems might contribute to the pathophysiology of type 2 diabetes, it will also be important to understand the regulation of these neurons in diabetic states, as well as in obesity. Even absent a comprehensive understanding of their regulation, however, VMN^Cckbr^ neurons could represent a pharmacologic target by which to improve glycemic control in all forms of diabetes.

## Methods

### Animals

Mice were bred in our colony in the Unit for Laboratory Animal Medicine at the University of Michigan; these mice and the procedures performed were approved by the University of Michigan Committee on the Use and Care of Animals (Protocol# 00011066) and in accordance with Association for the Assessment and Approval of Laboratory Animal Care and National Institutes of Health guidelines. Mice were provided with food and water *ad libitum* (except as noted below) in temperature-controlled rooms on a 12-hour light-dark cycle. For all studies, animals were processed in the order of their ear tag number, which was randomly assigned at the time of tailing (before genotyping). ARRIVE guidelines were followed; animals were group-housed, unless otherwise noted.

*Cckbr^Cre^* mice were previously described^15^. Prior to experiments, mice were genotyped via PCR across the genomic region of interest. Animals were processed and studied in the order of their randomly assigned ear tag number and investigators were blinded to genotype and/or treatment for all studies. All experiments were conducted with approximately equal number of males and females unless otherwise indicated.

### Reagents

Viral reagents AAV-DIO-ChR2-eYFP, AAV-DIO-GFP, and AAV-DIO-TT were as previously described^15^ and were packaged by the University of Michigan Viral Vector Core (University of Michigan, Ann Arbor, Michigan, USA).

### Stereotaxic injections

Eight- to 12-week-old animals were anesthetized with 1.5% to 2% isoflurane in preparation for craniotomy. After exposing the skull, bregma and lambda were leveled, a hole was drilled, the tip of a pulled glass pipette was inserted at the coordinates of our target, and the contents of the pipette were released at approximately 25 nL/min. For the VMN, 100 - 200 nL virus was injected at anteroposterior (AP) –0.750, mediolateral (ML) ±0.35, and dorsoventral (DV) –5.60. After 3 minutes to allow the virus to diffuse into the brain, the pipette was raised slowly from the hole in the skull, the hole in the skull was filled with bone wax, and the skin was closed with surgical sutures (Henry Schein). Analgesics were administered prophylactically to all mice to prevent postsurgical pain. Mice were allowed 4 weeks to recover from surgery before experimental manipulation. Analysis of fluorescent reporter expression was used to confirm proper targeting of the brain region in all experiments. Cases lacking expression in the region of interest were omitted from analyses.

### Optogenetics

Optogenetic fibers were placed 0.5 mm above the injection site for the VMN (AP: -0.75 mm, ML:0.35 mm, and DV: -5.10 mm from bregma) and were anchored above the skull with Metabond. Mice were allowed at least 4 weeks to recover from surgery before stimulation experiments. Laser stimulation (473nm, MBL-F-473, Opto Engine) was performed at 5-ms pulses, 40 pulses per second for a total of 30 – 60 minutes with an approximate irradiance 21 mW/mm^2^. Parameters were chosen to match previous optogenetic VMN stimulation that examined glucose mobilization^15,25^. Mice were fasted for 2 hours in the early light period prior to optogenetic stimulation. Blood was obtained from the tail vein and blood glucose was measured with Accu-Chek blood glucose meter (Accu-Chek Aviva Plus, Roche Diagnostics) for data from Fig 1A, and all glucose measurements in Figure 2. Blood glucose for figure 1C was measured in 10uL whole blood lysed in 1mL RUO Glucose/Lactate Hemolyzing Solution (EKF Diagnostics) using a Biosen C-Line Glucose and Lactate Analyzer (EKF Diagnostics). Blood for gluconeogenic substrate and hormone measurement was collected in EDTA-coated microtubes.

### Hormone and metabolite measurements

Insulin and glucagon were measured according to manufacturer’s instructions (Crystal Chem Ultra High Sensitive Mouse Insulin ELISA, cat: 90080; Mercordia Glucagon ELISA, cat: 10-1281-01). Corticosterone was measured in 20μL of serum by liquid chromatography-tandem mass spectrometry as previously described^37^. Plasma pyruvate, amino acids, glycerol, and non-esterified fatty acids were measured with a commercial colorimetric kit according to manufacturer’s instructions as follows: pyruvate (Sigma Aldrich, MAK071, 10μL), amino acids (Abcam, ab65347, 5μL), glycerol (Sigma Aldrich, MAK117, 10 μL), NEFA (Sigma Aldrich MAK044, 5 μL). All assays were performed in duplicate and compared to the provided standard curve. Lactate was measured in 10uL whole blood lysed in 1mL RUO Glucose/Lactate Hemolyzing Solution (EKF Diagnostics) using a Biosen C-Line Glucose and Lactate Analyzer (EKF Diagnostics). Plasma ketone levels were measured using 1.5μL of tail blood on Precision Xtra Blood Ketone Test Strips and a handheld Precision Xtra Blood Glucose and Ketone Meter (Abbott Labs). For liver glycogen measurements, liver was collected following euthanasia with isoflurane, frozen on dry ice and stored at -80C. For liver homogenization, 10 – 20mg of frozen liver was homogenized in 100uL sterile water, boiled for 5 minutes, and centrifuged at 13,000rpm for 5 minutes. The supernatant was then diluted 1:1 with sterile water and 20μL was used for glycogen measurement (Glycogen Assay Kit, Cat: MAK016, Sigma).

### Perfusion and IHC to confirm viral targeting

To confirm viral targeting for all mice, brains were collected and processed as previously described^17^. Briefly, mice were anesthetized with isoflurane before transcardial perfusion with PBS followed by 10% formalin. Brains were removed and placed into 10% formalin overnight, followed by 30% sucrose for at least 36 hours. Brains were cut into 30-μm sections on a freezing microtome in 4 series and stored in anti-freeze solution (25% ethylene glycol, 25% glycerol). Sections were washed with PBS and then pretreated for 1 hour in blocking solution (PBS containing 0.1% triton, 3% normal donkey serum) and then incubated overnight in blocking solution containing chicken anti-GFP (1:1000, GFP-1020, Aves). The following day, sections were washed in PBS, treated with blocking solution containing anti-chicken Alexa Fluor 488 fluorescent secondary antibody (1:200), and washed again in PBS. Sections were mounted onto slides and cover-slipped with Fluoromount-G (Southern Biotech). Slides were imaged using an Olympus Bx53 microscope.

### Phenotypic studies

For glycemic tests, mice were fasted for 4 hours starting at lights on. Mice were injected with glucagon (200 μg/kg, i.p.; Sigma, St Louis, MO, USA), pyruvate (2g/kg, i.p; Sigma, St Louis, MO, USA), or glycerol (2g/kg, i.p.; Sigma, St Louis, MO, USA). Blood was obtained from the tail vein and blood glucose was measured with Accu-Chek blood glucose meter (Accu-Chek Aviva Plus, Roche Diagnostics). For optogenetic stimulation following glycogenolysis inhibitor CP91,149 (25mg/kg unit/kg, oral gavage, Cayman Chemicals, Ann Arbor, MI, USA), CP91,149 was prepared as follows: CP91,149 was resuspended in DMSO at 25mg/mL. 10.9mL of 1% methylcellulose (Sigma) in sterile water was added to 100μL CP91,149, then 2μL Tween-80 was added the solution vortexed until evenly suspended. Vehicle was prepared exactly as CP91,149 with DMSO. Mice were pretreated with CP91,149 or vehicle for 1 hour before optogenetic stimulation. For optogenetic stimulation following β3-receptor antagonist SR-59230A, mice were injected with SR-59230A (1.5 unit/kg, i.p., Sigma) 2 hours prior to optogenetic stimulation.

### Glycerol Appearance Assay

Glycerol appearance assays were performed with the assistance of the University of Michigan Animal Phenotyping Core. Under anesthesia, mice were catheterized with silicon tubing. After surgery, mice were housed individually, and their body weight was monitored daily. Mice that lost more than 10% of their presurgery body weight were removed from the study. On the day of data collection, mice were fasted for 2.5 hours starting 3 hours after lights-on. [2-^3^H]glycerol was infused using a bolus-continuous infusion protocol of 2μCi in 2min + 0.1μCi/min for a 90-minute equilibration period. After the equilibration period, [2-^3^H]glycerol was infused at a steady state and plasma collected for blood glucose, plasma glycerol and [^3^H] plasma measurements as previously described^38^. Thirty minutes after initial plasma collection, β3-agonist CL-316,243 (1.5mg/kg, i.v. Sigma) was injected and plasma glucose, glycerol, and plasma [^3^H] radioactivity were measured at baseline and after 10 minutes.

### Continuous Glucose Monitoring

Continuous glucose monitors were surgically implanted, and data was collected with assistance from the University of Michigan Physiology Phenotyping Core. Briefly, under isoflurane anesthesia mice were shaved on the ventral neck and a midline skin incision from chin to mid-sternum was performed, and the salivary glands were retracted. The left common carotid artery was dissected and distally ligated and proximally occluded with silk sutures. A bent 25-gauge needle was inserted into the common carotid and the glucose sensor tip of a HD-XG mouse glucose telemetry implant (Data Sciences International, St. Paul, MN) was inserted and then advanced to the aortic arch and secured with tissue adhesive and a silk suture. The body of the device was rotated caudally and inserted into a subcutaneous pocket that was created over the right ventral flank. The glands were reapproximated and the skin incision closed with 2-3 staples. The animal was returned to its housing cage and placed on a heating pad until it was mobile.

Following surgical recovery, singly housed mice were placed on a receiver plate. To calibrate the glucose telemetry probe, a tail vein blood sample was taken to measure glucose and the value was paired with the corresponding voltage reading such that the software could give glucose measurements.

### RNA Quantification

To determine liver mRNA expression, liver was collected and frozen on dry ice. Liver tissue was homogenized and mRNA was then extracted using the RNeasy Kit (Cat #74104, Qiagen) according to the manufacturer’s instructions; cDNA was subsequently produced using the iScript cDNA synthesis kit (BioRad). Real-time quantitative PCR was conducted using a StepOnePlus Real-Time System (Applied Biosystems) with Radiant Probe Hi-ROX qPCR Kit (Alkali Scientific) and Taqman Gene Expression Assays (Thermo Fisher Scientific). The expression of target genes was normalized to *Gapdh* expression.

### Adipocyte Quantification

Adipocyte size was quantified for gonadal and posterior subcutaneous white adipose depots that were sectioned and stained with hematoxylin and eosin. Adipocyte cross-sectional areas were calculated using Adiposoft (Version 1.15) plug-in in Fiji, or ImageJ (Version 2.9.0). Average adipocyte size per group was calculated by comparing the averages of each sample between groups. Differences in means were statistically analyzed using an unpaired t-test (p<0.05) in GraphPad Prism 9.0 (Version 9.3.1). Frequency distribution curves were generated for each test group using bin range: 0 to 8,000 μm^2^ and bin width: 250 μm^2^. Curves were fit to the data using a non-linear, lognormal Gaussian regression in GraphPad Prism 9.0 (Version 9.3.1)^22^.

### Statistical Analysis

Sample sizes were not predetermined using statistical tests but were similar to previous studies with similar approaches^15^. Figure 1 and Figure 2 C-D were analyzed using a paired Student’s *t* test. Figure 2A, Figure 3 B-E, Figure 4, Figure 5, and Figure 6 data were analyzed by unpaired Student’s t test. Data were omitted if the injection missed or the animal was sick or injured at the time of the experiment (loss of >10% BW). Continuous glucose monitoring was separated into 4-hr circadian windows and each 4hr window was separately analyzed by 2-way ANOVA. Significance was set at a P value of 0.05 or less. All data analysis was performed using GraphPad Prism software (GraphPad Software).

## Author Contributions

AH, NK, JS, AJT, WTY, JNF, and AHA performed most experiments and analyzed and interpreted data. KTL, LRO, and HM performed WAT depot and adipocyte measurements, data analysis and interpretation. NBK performed plasma hormone measurements and data analysis. AFT, OAM, MGM and AHA designed and supervised the studies, analyzed and interpreted data. AH and AHA wrote the manuscript, with editing by MGM. All authors discussed the results and commented on the paper.

## Acknowledgements

Research support was provided by the Michigan Diabetes Research Center (NIH grant P30 DK020572), the Mouse Metabolic Phenotyping Center – Live (U2CDK135066) Physiology Phenotyping Core, the Michigan Nutrition and Obesity Center Adipose Tissue Core (P30 DK089503); Department of Veterans Affairs (IK2BX005715 to NBK); the Warren Alpert Foundation (Warren Alpert Distinguished Scholars Award to AHA); Endocrine Fellows Foundation (EFF Research Grant to AHA), the Marilyn H. Vincent Foundation (to MGM); and Novo Nordisk (to MGM). This work was also supported in part by NIH grants K08 DK1297226 (to AHA). We thank Bjoern Brixius for assistance with mass spectrometry measurement of corticosterone.

